# Non-trivial compositionality in chimpanzees: an experimental demonstration

**DOI:** 10.64898/2026.10.05.756726

**Authors:** Aurélie Le Fur, Alban Lemasson, Simon W. Townsend, Maël M. Leroux

## Abstract

Compositional syntax—the ability to combine meaningful units into structured sequences whose overall meaning depends on both the meanings of the parts and their combination—is a hallmark of human language. Recent work suggests that chimpanzees may exhibit similar capacities, raising the possibility that compositionality predates the emergence of language in humans. However, a key challenge has been to exclude simpler explanations in which call combinations are processed as independent utterances rather than integrated structures.

Here, we address this issue using a source-manipulation paradigm in playback experiments with captive chimpanzees (*n* = 6) at the Sigean African Reserve, France. We presented sequences combining alarm and recruitment calls either from a single source (one loudspeaker) or from two spatially distinct sources (two loudspeakers). If sequences are interpreted as independent calls, responses should not differ between conditions. Instead, chimpanzees responded significantly faster when sequences originated from a single source. Bayesian survival analyses confirmed this effect.

These results support a non-trivial form of compositionality, whereby the meaning of call combinations depends on their integration into a unified sequence produced by a single individual. Our findings provide experimental evidence that key features of linguistic compositionality may have evolutionary roots in the primate lineage.

## Introduction

The capacity to combine discrete vocal elements into structured, meaningful sequences is central to language ^1^. Yet, a growing number of studies report that animals, including birds ^2,3^ and primates ^4,5^, also produce meaningful vocal sequences. In chimpanzees, for example, “alarm-huus” (AH) – emitted in response to danger – and “waa-barks” (WB) – used for recruitment – are combined into AH–WB sequences during snake encounters ^6^. These sequences have been characterized as compositional, since the meaning of the whole is derived from its parts ^7^. Whilst intriguing and representing striking parallels with human language, a recurring critique is that rather than being truly compositional, such structures may rather represent two separate utterances ^8^. In other words, it remains unclear whether such a sequence constitutes a single syntactic construction produced by the signaler and interpreted as a single utterance by the receiver (non-trivial compositionality), or whether the sequence is in fact two separate utterances whose meanings are combined by the receiver a posteriori (trivial compositionality) ^8,9^. Disentangling these two scenarios in apes is absolutely central to reconstructing the evolutionary progression of human compositionality, yet experimental demonstration of non-trivial compositionality in our closest living relatives is still lacking. To address this, we adapted a method originally developed for birds ^10^, in which calls from a combination are broadcast either from a single source (one loudspeaker mimicking a situation in which the combination is produced by a single individual) or from two separate sources (two loudspeakers mimicking a situation in which the two calls are produced by two distinct individuals) (see Figure 1). This allowed us to test whether compositional processing is contingent on the combination being produced as a single, bona fide signal. If responses were to differ between the two conditions, this would indicate that the combination is non-trivial: the two calls have to be linked structurally, i.e. produced by a single individual, for the combination to be meaningful for the receiver. Conversely, if chimpanzees were to respond similarly in both conditions, this would suggest that receivers can interpret the combination regardless of whether it is produced as a single utterance or as two separate utterances, indicating a case of trivial compositionality.

**Figure 1.**
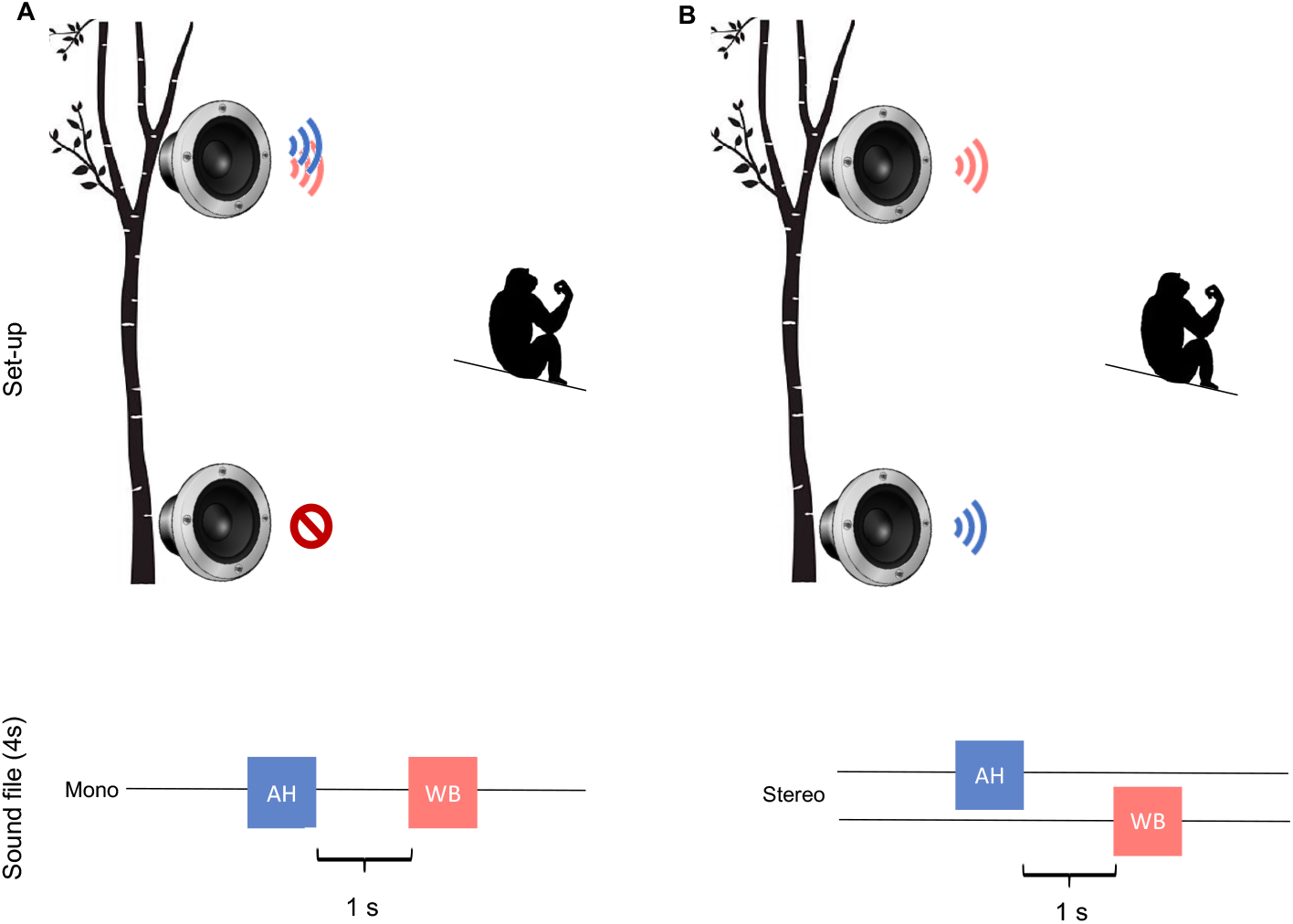
Setup of the experiment (inspired by ^10^). **A:** 1-Individual condition with the active loudspeaker in the top position, **B:** 2-individuals condition with loudspeaker broadcasting the “Alarm-huu” in the bottom position. AH = “Alarm-huu”, WB = “Waa-bark”.

We tested six adult chimpanzees housed at the Réserve Africaine de Sigean, France. Previous observations showed that the “alarm-huu” and “waa-bark” combination is part of the vocal repertoire of this group, confirming that our stimuli reflected naturalistic behavior ^11^. Subjects were exposed to playback stimuli consisting of one “alarm-huu” followed by one “waa-bark”, separated by a one-second interval ^6^. In the 1-individual condition, both calls were played from a single loudspeaker, mimicking a single caller. In the 2-individuals condition, the calls were played from two spatially distinct loudspeakers (2.5 m apart, one on the ground, one in a tree, see Figure 1), mimicking a combination originating from two different callers, albeit communicating potentially about a single event. Each subject experienced both conditions, with playback order and loudspeaker positions randomized. The playback stimuli were taken from unfamiliar wild chimpanzees recorded during snake encounters ^6^. We measured three behavioral responses: (i) latency to first look towards the loudspeakers (calculated from the onset of the second call), (ii) total gaze duration within two minutes, and (iii) the number of looks ^6^. We analyzed (i) and (ii) using survival Bayesian models (significant if the 95% Credible Interval [CI95%] does not cross 1) and (iii) using a Bayesian negative binomial model (significant if CI95% does not cross 0). We controlled for potential confounding factors. First, to account for social facilitation (i.e. the possibility that the subject’s response was influenced by the reaction of other group members), the proportion of individuals reacting was included as a fixed effect. Second, to control for potential habituation across trials, we included trial order as a random effect. To further reduce habituation, we alternated between two loudspeaker locations, and we tested the effect of this variable by including it as a random effect in our models. Finally, since each individual were tested in both conditions, we also fitted identity as a random effect.

## Results

Chimpanzees responded significantly faster in the 1-individual condition than in the 2-individuals condition (Hazard Ratio = 0.04, CI_95%_ = [0.003, 0.26]), indicating a shorter latency to look when the sequence appeared to come from a single individual (Figure 2). There was no significant effect of condition on gaze duration (HR = 0.73, CI_95%_ = [0.22, 2.51]) or number of looks (β = –0.48, CI_95%_ = [–1.61, 0.62]). The proportion of group members reacting to the playback had no effect on any measure (latency: HR = 0.71, CI_95%_ = [0.20, 2.29]; gaze duration: HR = 0.65, CI_95%_ = [0.30, 1.43]; number of looks: β = 0.30, CI_95%_ = [–0.42, 0.98]). Trial order, loudspeaker location, and subject identity did not significantly influence responses (1∈CI_95%_ for latency and gaze duration and 0∈CI_95%_ for number of looks for all variables). These results suggest that, although chimpanzees did not look longer or more often at the loudspeaker across conditions, they were quicker to look when a sequence was originating from a single loudspeaker.

**Figure 2.**
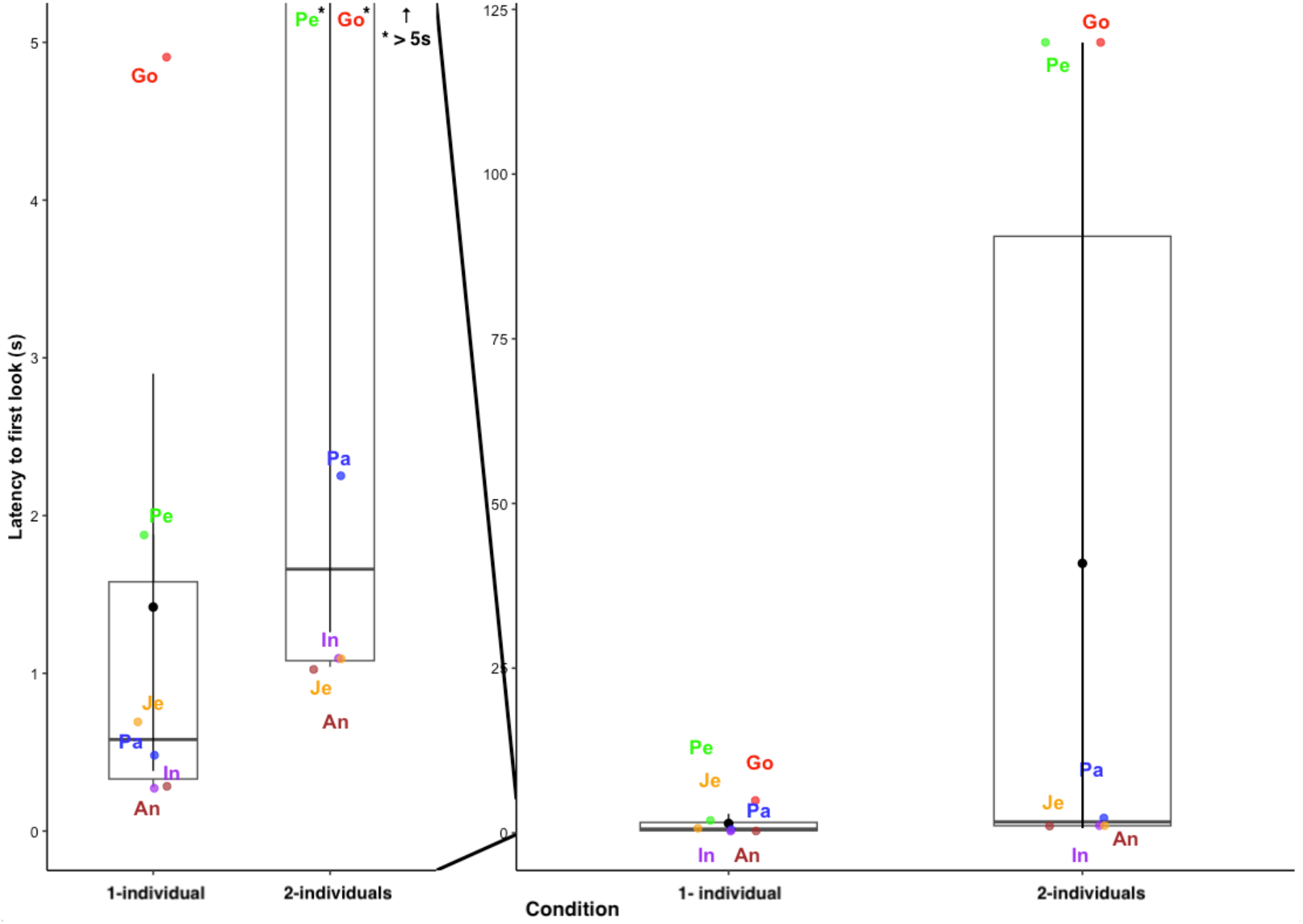
Influence of the condition (1-individual or 2-individuals) on the latency to first look towards the loudspeakers. Individuals are represented with their color code and the first two letters of their name. The plot on the left shows a zoom on latencies between 0 and 5 seconds (* Pe and Go have a latency superior to 5 seconds in the 2-individuals condition).

## Discussion

Results show that when both calls from the combination were presented from a single source, chimpanzees reacted faster, indicating that the sequence carried greater urgency, as would be expected when a single individual simultaneously signals the presence of a snake and the need for immediate assistance. In contrast, when presented from two distinct sources, the combination elicited slower reactions from chimpanzees. Here, individuals seemed to process the combination as separate utterances, where integrating the meaning of both calls may require additional inferential processing from the individuals. In sum, this indicates that in the 1-individual condition, chimpanzees likely treated the sequence as a single, compositional utterance conveying an urgent message. Such findings provide experimental support for non-trivial compositionality in chimpanzees, where receivers seem to not only attend to the semantic content of each call in a sequence independently, but also to the precise way in which they are combined.

The implications of these findings are significant for our understanding of language evolution. Distinguishing trivial from non-trivial compositionality is essential when evaluating claims of “syntax” in non-human species. Not all combinations qualify as syntactic or compositional, and experimental designs that manipulate the number of emitters or sequence integrity can reveal whether animals are interpreting a structure holistically, a key feature of human syntax. More specifically, our experimental findings complement recent observational work indicating non-trivial compositionality in the bonobo vocal system ^5^. The presence of non-trivial compositionality in both bonobos and now chimpanzees suggests that this core feature of language is more common than previously thought and may even have evolved since the last common ancestor of Homo and Pan.

## Materials and Methods

### Study Site and Subjects

Data were collected at the Réserve Africaine de Sigean (RAS), a 350-hectare zoological park in southern France housing a single group of 11 chimpanzees (*Pan troglodytes*). The group has remained stable for over 13 years. The chimpanzees had access to two connected forested islands (1.5 ha total) and a 90 m^2^ indoor night lodge. Outdoor access was granted daily. Environmental enrichment (ropes, burlap, artificial termite mounds, platforms) was provided in both areas. Individuals were fed six times per day and had ad libitum access to water. The study involved six adults (4 females, 2 males; aged 14–51 years). All procedures complied with EU Directive 2010/63/EU and were approved by the University of Rennes Animal Ethics Committee (Ref: #2024031015037725).

### Playback Stimuli

We generated 12 unique stimuli (1 per subject per condition), using alarm vocalizations from wild chimpanzees recorded in the Sonso community (Budongo Forest, Uganda) during snake presentations ^6^. Each stimulus combined one “alarm-huu” and one “waa-bark” separated by a 1s silence ^6^. In the 1-individual condition, both calls originated from the same male; in the 2-individuals condition, calls were produced by different males. All recordings were edited using Audacity 2.4.2 (.wav, 44.1 kHz, 32-bit), normalized to 23 dB (RMS criterion), and faded in and out to minimize acoustic artefacts. The mean duration of the stimuli did not differ between conditions (i.e. 2.886 ± 0.96s). Only male vocalizations were used to match prior studies and avoid confounding sex effects.

### Experimental Procedure

To test chimpanzees’ sensitivity to non-trivial compositional structuring, we adapted the design of Suzuki & Matsumoto ^10^, who simulated sequences from one or two individuals using spatially separated loudspeakers. To avoid attentional biases arising from horizontal displacement in the original setup, we implemented a vertical loudspeaker configuration. Two camouflaged loudspeakers were placed 2.5 m apart: one on the ground and one ∼2 m high in a tree. Both loudspeakers were present in all trials, but only one was active in the 1-individual condition. The active loudspeaker position was randomized in this condition. In the 2-individuals condition, the “alarm-huu” and “waa-bark” were emitted from different loudspeakers. The loudspeaker emitting the first call was randomized in this condition, and the order of conditions was randomized across trials.

In accordance with ethical and welfare considerations, all subjects were tested while remaining in their social group, and no individuals were isolated for the purposes of the experiment. Playback occurred when the focal subject had been resting or socializing calmly for at least 2 min, with no recent disturbance (e.g., vocalizations, agonism). Stimuli were played when the subject was not facing the loudspeakers to allow gaze shifts to be measured. Each subject experienced one trial per condition. Trials were separated by ≥24 h (on average separated by 5 days ± 3 days) and occurred at different times and locations to minimize habituation. Two fixed external spots (A and B) were used to simulate the presence of outgroup individuals.

Responses were filmed using three video cameras: a handheld Canon LEGRIA HF R806 to focus on the subject; a GoPro HERO10 placed on a balcony for an overhead view; and a JVC GZ-R435BEU positioned to film other group members. Filming began 5 min before playback and continued until the subject returned to baseline or for at least 5 min post-playback. The experimenter remained in areas of the enclosure where the chimpanzees were accustomed to seeing them and never looked towards the loudspeakers to limit experimenter effects.

### Behavioral Coding and Statistical Analysis

Videos were coded frame-by-frame using BORIS v8.24.1 ^12^. Following Leroux et al. ^6^, we coded three variables: the number of looks toward the loudspeakers, the total gaze duration within 2 min post-playback, and the latency to first look calculated from the onset of the second call. A latency of 120 s was assigned if the subject did not look. Coding reliability was assessed by a second observer on 33% of trials; intra-class correlation coefficients for all measures exceeded 0.8.

Data were analyzed in R v4.4.0 using the brms package ^13^. We modelled the number of looks using a Bayesian negative binomial regression. We interpreted fixed effects based on their 95% credible intervals (CIs); if the CI crossed zero, we considered that there was no clear evidence for an effect. Latency and duration were analyzed using survival analyses, allowing for the inclusion of trials in which there was no response, through Bayesian Cox proportional hazards models. We used the hazard ratio to interpret fixed effects. It indicates the relative rate at which a behavior occurs. If the 95% CI includes 1.00, we considered there was no clear difference between effects. All models included subject identity, trial order, and location (spot A or B) as random effects. The proportion of group members visibly reacting (e.g., looking, moving, or vocalizing) was included as a fixed effect to control for social facilitation. For the number of looks, the initial model without priors failed to converge. Convergence was achieved using informative priors: normal (1,1) for intercepts, normal (0,1) for fixed effects, student_t (3,0,2.5) for standard deviations of random effects, and gamma (2,0.1) for shape parameters. All models showed satisfactory convergence (Ř < 1.01).

## Ethical Note

All procedures complied with EU Directive 2010/63/EU and were approved by the University of Rennes Animal Ethics Committee (Ref: #2024031015037725).

## Data, Materials, and Software Availability

Raw data and analysis scripts are available as supporting information. Supplementary tables and results are available on the Open Science Framework.

## Acknowledgments

We thank Lyna Rachid-Martin, Axelle Verges, Antoine Joris and Marielle Beltrame and the staff of the Réserve Africaine de Sigean for their support. We thank Emmanuel Chemla and Philippe Schlenker for useful discussions. S.W.T. was supported by the Swiss National Science Foundation (grant ID: SNSF 315130_192620, SNSF PP00P3_163850 and SNSF PP00P3_198912). M.M.L. was supported by the French National Agency for Research and Rennes Metropole (grant IDs: ANR-24-CE28-6179-01 and 24CO981).

