## Supplementary methods for "Non-trivial compositionality in chimpanzees: an experimental demonstration"

**This PDF file includes:**

Supporting Results

**Supporting Information Text**

**Supplemental Results**

We found no significant effect of condition (1-individual vs. 2-individuals) on total gaze duration toward the loudspeakers (Hazard Ratio [HR] = 0.73, $CI_{95\%}$ = [0.22, 2.51]) or number of looks (β = –0.48, $CI_{95\%}$= [–1.61, 0.62]) (Tables S2–S3). However, chimpanzees looked significantly faster in the 1-individual condition than in the 2-individuals condition (HR = 0.04, $CI_{95\%}$ = [0.003, 0.26]; Table S2), indicating a shorter latency to orient toward the sound source when the call sequence originated from a single individual.

The proportion of other individuals visibly reacting to the playback had no significant effect on any of the response variables (gaze duration: HR = 0.65, $CI_{95\%}$ = [0.30, 1.43]; number of looks: β = 0.30, $CI_{95\%}$ = [–0.42, 0.98]; latency: HR = 0.71, $CI_{95\%}$ = [0.20, 2.29]). Random effects for trial order and location had negligible influence on model outcomes (see Tables S4–S6 for full model summaries).

**Table S1:** Hazard ratio for Bayesian survival models comparing behavioural responses (gaze duration and latency to first look) relative to the tested condition (1-individual or 2-individuals) and the proportion of individuals who have reacted. A hazard ratio gives the relative rate at which a behaviour of interest occurs; if the lower and upper bounds overlap with 1.00, it indicates a lack of reliable difference between the effects. The effect with a $CI_{95\%}$ that does not include 1 is shown in bold.

| **Behavioural response** | **Fixed effect** | **Median** | **SD** | **Hazard ratio lower 95% bound** | **Hazard ratio upper 95% bound** |
|---|---|---|---|---|---|
| **Gaze duration** | Condition | 0.730 | 0.954 | 0.223 | 2.510 |
| | Proportion of individuals who reacted | 0.647 | 0.421 | 0.299 | 1.430 |
| **Latency to first look** | **Condition** | **0.037** | **0.503** | **0.003** | **0.259** |
| | Proportion of individuals who reacted | 0.710 | 0.872 | 0.197 | 2.290 |

**Table S2:** Results of the Bayesian negative binomial model testing the influence of the condition (1-individual/2-individuals) and the proportion of individuals who have reacted on the number of looks.

| Predictor | Estimate | SD | 95% CI Lower | 95% CI Upper |
|---|---|---|---|---|
| Condition | -0.475 | 0.560 | -1.609 | 0.616 |
| Proportion of individuals who reacted | 0.298 | 0.344 | -0.415 | 0.976 |

**Table S3:** Results of the Bayesian survival model testing the influence of the condition (1-individual/2-individuals) on the gaze duration.

| Effect Type | Predictor | Estimate | SD | 95% CI Lower | 95% CI Upper |
|---|---|---|---|---|---|
| Fixed Effect | Intercept | 1.430 | 1.336 | -1.054 | 4.512 |
| Fixed Effect | Condition | -0.302 | 0.756 | -1.771 | 1.193 |
| Fixed Effect | Proportion of individuals who reacted | -0.427 | 0.498 | -1.386 | 0.600 |
| Random Effect | Identity | 0.745 | 0.664 | 0.028 | 2.453 |
| Random Effect | Order | 0.735 | 0.593 | 0.033 | 2.216 |
| Random Effect | Spot | 1.372 | 1.303 | 0.038 | 4.728 |

**Table S4:** Results of the Bayesian survival model testing the influence of the condition (1-individual/2-individuals) on the latency to first look.

| Effect Type | Predictor | Estimate | SD | 95% CI Lower | 95% CI Upper |
|---|---|---|---|---|---|
| Fixed Effect | Intercept | 3.870 | 1.690 | 0.308 | 7.250 |
| Fixed Effect | Condition | -3.388 | 1.426 | -6.431 | -0.795 |
| Fixed Effect | Proportion of individuals who reacted | -0.369 | 0.792 | -2.044 | 1.080 |
| Random Effect | Identity | 1.747 | 1.029 | 0.136 | 4.218 |
| Random Effect | Order | 1.195 | 0.916 | 0.049 | 3.418 |
| Random Effect | Spot | 1.880 | 1.608 | 0.078 | 5.962 |

**Table S5:** Results of the Bayesian negative binomial model testing the influence of the condition (1-individual/2-individuals) on the number of looks.

| Effect Type | Predictor | Estimate | SD | 95% CI Lower | 95% CI Upper |
|---|---|---|---|---|---|
| Fixed Effect | Intercept | 1.492 | 0.732 | -0.009 | 2.912 |
| Fixed Effect | Condition | -0.475 | 0.560 | -1.609 | 0.616 |
| Fixed Effect | Proportion of individuals who reacted | 0.298 | 0.344 | -0.415 | 0.976 |
| Random Effect | Identity | 0.735 | 0.533 | 0.040 | 2.077 |
| Random Effect | Order | 0.655 | 0.482 | 0.032 | 1.838 |
| Random Effect | Spot | 1.372 | 1.160 | 0.100 | 4.395 |
